# Reaction pathways for the enzymatic degradation of poly(ethylene terephthalate): What characterizes an efficient PET-hydrolase?

**DOI:** 10.1101/2022.09.13.507771

**Authors:** Sune Schubert, Kay Schaller, Jenny Arnling Bååth, Cameron Hunt, Kim Borch, Kenneth Jensen, Jesper Brask, Peter Westh

## Abstract

Bioprocessing of polyester waste has emerged as a promising tool in the quest for a cyclic plastic economy. One key step is the enzymatic breakdown of the polymer, and this entails a complicated pathway with substrates, intermediates, and products of variable size and solubility. We have elucidated this pathway for poly(ethylene terephthalate) (PET) and four enzymes. Specifically, we combined different kinetic measurements and a novel stochastic model, and found that the ability to hydrolyze internal bonds in the polymer (endo-lytic activity) was a key parameter for overall enzyme performance. Endo-lytic activity promoted the release of soluble PET fragments with two or three aromatic rings, which, in turn, were broken down with remarkable efficiency (k_cat_/K_M_-values of about 10^5^ M^-1^s^-1^) in the aqueous bulk. This meant that about 70% of the final, monoaromatic products was formed via soluble di-or tri-aromatic intermediates.

## Introduction

Bioprocessing of plastic waste streams is a promising tool in the battle against environmental problems associated with the escalating consumption of plastics ^1^. Currently, the most developed technology in this area is monomer recovery after enzymatic hydrolysis of the widely used polyester poly(ethylene terephthalate), PET^2–5^. PET-degrading enzymes (henceforth called PET-hydrolases) have been known for decades^6^ and although some specificity for PET has been identified among different families of hydrolases, effective degraders of bulk PET with application potential have typically been found among enzymes classified as cutinases [EC 3.1.1.74]^7^. Recently, PET-hydrolases with significantly improved performance have been discovered^8–10^ or engineered^11–13^, and this has spurred optimism for full-scale PET bioprocessing. The hydrolysis of all ester bonds in a PET chain would release the two monomeric products, terephthalic acid (T) and ethylene glycol (E), but many studies have reported a dominance of soluble products with one or two conserved ester bonds. Soluble PET fragments include mono-(2-hydroxyethyl)terephthalate (henceforth abbreviated ET) and bis(2-hydroxyethyl) terephthalate (abbreviated ETE)^1^, which tend to accumulate in hydrolyzates because PET-hydrolases typically show poor activity against these molecules^14^.

Despite the progress mentioned above, implementation of PET bioprocessing will probably require continued optimization of the enzymatic process. Some key goals in this process include discovery or engineering of better industrial enzymes, improved methods for substrate pretreatment, and a better fundamental understanding of the reaction course and hence bottlenecks for the degradation of the polymer^15,16^. In the current work, we address the latter of these tasks, by analyzing the pathway from insoluble polymer to soluble monomers. This pathway is complex because all enzyme products, except the monomers T and E, are potential substrates for further esterolytic reactions. This aspect has been discussed previously by Zimmermann and co-workers, who monitored PET hydrolysis by a range of analytical techniques, and concluded that the reaction progressed as a combination of endo- and exo-lytic bond cleavage^17,18^. Endo-lytic cleavage (i.e., hydrolysis of an ester bond located internally in the polymer) may not release soluble product (except for a proton), but modifies the reactivity of the PET surface. Conversely, exo-lytic reactions (bond cleavage near the polymer end) release a soluble product, but probably generates little or no change in the reactivity of the remaining polymer. PET fragments with two or three aromatic moieties show some aqueous solubility (see below), and it follows that reaction products of this type will transfer to the bulk phase, where they become yet another type of substrate. All in all, this generates a complicated network of possible reaction pathways between polymeric PET and the monomeric products, and routes through this network will depend on the enzyme’s specificity for substrates with different physical properties.

In an attempt to unveil examples of these reaction pathways, we have used complementary experimental methods to monitor the enzymatic activity on both insoluble PET (nanoPET) and soluble PET fragments with 1, 2, or 3 aromatic moieties. Soluble substrates were characterized by steady-state kinetics, and for insoluble PET we monitored progress curves for the release of both protons and organic products. To systematically interpret the experimental data, we implemented a novel stochastic model for kinetic analyses of PET degradation. We found that the model was able to account for the experimental data and we used this to trace fluxes through different reaction pathways. Comparison of four PET-hydrolases showed differences between the highly efficient Leaf Compost Cutinase^19^ (LCC) and the PETase from *Ideonella sakariensis*^8^ (IsP) on one hand and less proficient PET-hydrolases from *Thermofibida fusca*^20^ (TfC) and *Humicola insolens*^18^ (HiC), on the other. We propose that these characteristic differences may help identify beneficial specificities of industrial PET-hydrolases.

## Results

We have characterized the activity of four PET-hydrolases by four complementary approaches. These were: i) progress curves for soluble, organic products ii) progress curves for proton release iii) steady-state kinetics for the hydrolysis of soluble PET fragments iv) stochastic modeling of progress curves.

### Progress curves on polymeric PET

We monitored the time course of PET degradation by both reversed-phase high-performance liquid chromatography (RP-HPLC) and pH-stat measurements. In the chromatographic analysis, we could resolve all three products with one aromatic moiety, i.e., T, ET, and ETE. In addition, we obtained peaks for products with two aromatic moieties, but we were unable to resolve the three different molecules, TET, ETET, and ETETE, in this group. The hydrolysates also showed a peak with a retention time corresponding to standards of the tri-aromatic fragment TETET, which was synthesized in house (see Experimental section). To further investigate tri-aromatic fragments, we also synthesized ETETETE, but the solubility of this (uncharged) molecule was below the detection level of the HPLC protocol. The anionic fragment TETET was more soluble and standard solutions showed a linear HPLC response up to at least 200 µM. Based on these observations and computed octanol-water partitioning coefficients^21^ for TETET, TETETE and ETETETE, we conclude that the HPLC peak for tri-aromatic fragments primarily reflected TETET and possibly some TETETE. Therefore, we propose, that TETETE provides a useful, operational boundary for soluble vs. insoluble products during PET hydrolysis.

Using the above RP-HPLC assignments, we calculated progress curves for five types of products (E, ET, ETE, di-aromatics (TET, ETET and ETETE combined), and tri-aromatics (TETET and ETETET combined), and plotted them in Fig. 1. The dominant species, T and ET, are given on the left ordinate, while the concentrations of the three other products are on the more expanded ordinate to the right.

**Figure 1.**
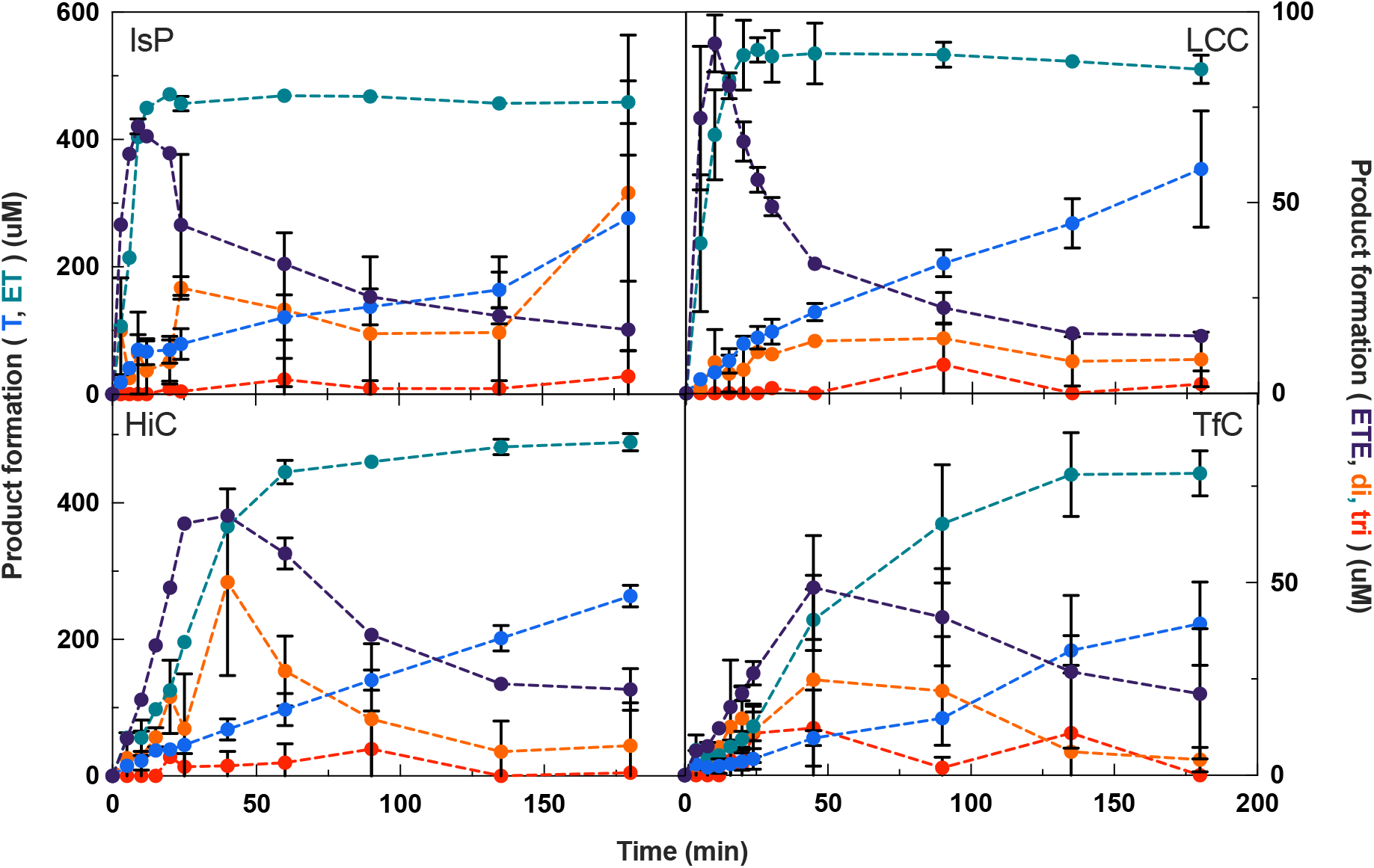
Progress curves for the degradation of nanoPET measured by HPLC. The chromatographic analysis separated five classes of products. These were the three mono-aromatic compounds T, ET, and ETE as well as accumulated amounts of products with two or three aromatic groups. The initial substrate load of nanoPET was 166.4 mg/L in all cases. For LCC, HiC, and TfC we used an enzyme concentration of 0.30 µM and an experimental temperature of 50 °C. The less thermostable IsP was assayed at 40 °C and 0.20 µM. Error bars represent the standard deviation of duplicate measurements.

ET was a dominant product for all four enzymes, but its release pattern fell into two groups. LCC and IsP (upper panels) showed substantial release of ET immediately after the reaction was started, while HiC and TfC (lower panels) displayed a lag-phase with sigmoidal ET-progress curves, which were steepest about 20 min into the process. This grouping of the enzymes was also visible for other products including ETE, which showed a distinct early peak at t∼10 min for LCC and IsP and subsequently disappeared gradually. For TfC and HiC, the peak in ETE was observed later at t = 30-50 min.

We monitored the real-time reaction progress in the same experiments by pH-stat measurements. Results in Fig. 2A showed that LCC and IsP released nearly 1 mM of protons within the first 10 min. Subsequently, these two enzymes showed very slow proton release. By contrast, HiC and TfC again revealed an initial lag phase and reached their maximal proton release rate after 15-25 min. As one proton is released for every ester bond cleaved, the results in the right ordinate of Fig. 2A provide a measure for the total degree of ester bond hydrolysis, D_hydrol_. Specifically, we may write;

**Figure 2.**
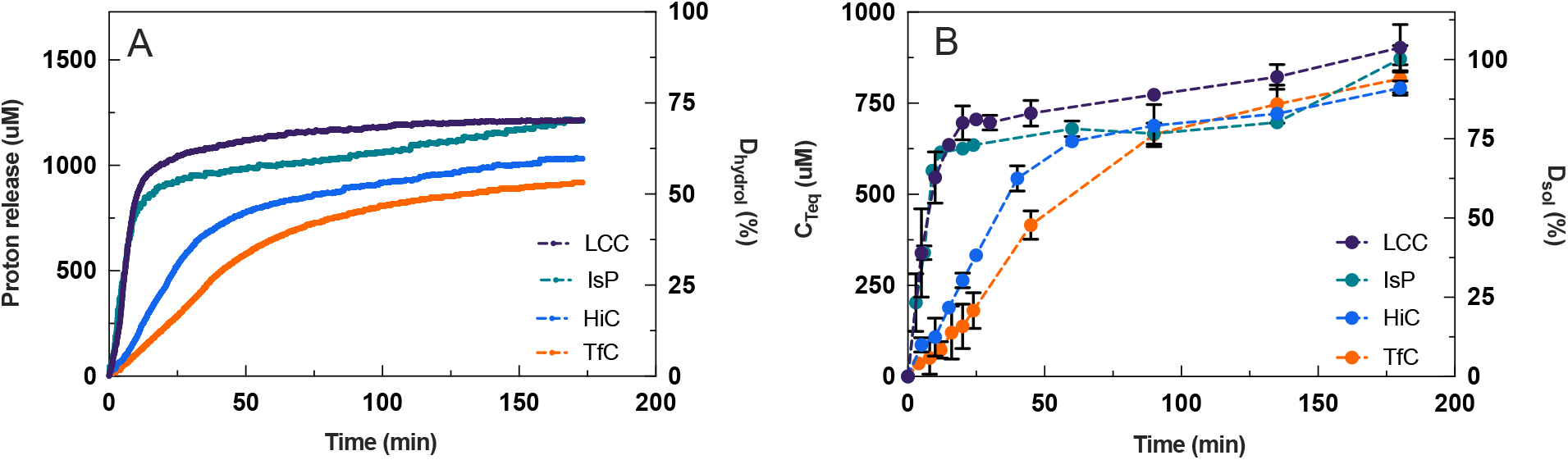
Progress curves for the hydrolysis of nanoPET by LCC, IsP, HiC, and TfC. Panels A and B show respectively the concentration of protons added in the pH-stat to keep the pH constant (pH 8.0) and the total concentration of soluble fragments in terephthalic acid equivalents, C^Teq^ (see Eq. 2). On the right ordinate, these functions are expressed in relative amounts as defined in Eqs. (1) and (3). Error bars represent the standard deviation of duplicate measurements.

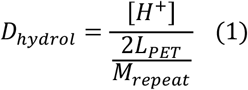

where [H^+^] is the concentration of protons specified on the left ordinate in Fig. 2A, L_PET_ is the initial load of PET (in g/L) and M_repeat_ is the molar mass of the repeating unit in PET (M_repeat_=192.2 g/mol). The factor of 2 in the denominator reflects that there are two ester bonds per repeating unit (neglecting end effects). This definition implies that if D_hydrol_ reaches 100%, the products will be stoichiometric amounts of T and E, while complete conversion of PET to ET, for example, would correspond to D_hydrol_ = 50%. The results in Fig. 2A (right ordinate) show that the aforementioned change from rapid to almost no proton release for LCC and IsP occurred for D_hydrol_ of 50-60%. It further appeared that the maximal rate for HiC and TfC (steepest progress curve for proton release) was observed for D_hydrol_ at around 20%.

It is of interest to directly compare the release of protons and organic products. For this purpose, we defined a total concentration of soluble products weighted in units of terephthalic acid (T) equivalents, C_teq_

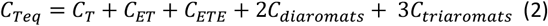

Symbols on the right side of Eq. (2) represent the concentrations of the five different species shown in Fig. 1. Plots of C_teq_ against time for the four enzymes are shown in Fig. 2B (left ordinate). It appeared that these progress curves followed basically the same overall course as the independently measured proton release data in Fig. 2A. However, at any given time, the concentration of protons (Fig. 2A) was larger than C_teq_ (Fig. 2B). This shift could reflect either endo-type cleavages, which release protons, but no other soluble product, or secondary hydrolysis of fragments in the bulk, which releases protons without affecting the number of T-equivalents in solution (*i*.*e* C_teq_ remains constant, *c*.*f*. Eq. 2). We will return to this in the discussion. First, we define a relative value of C_teq_. This parameter signifies the mole fraction of T-moieties that has been transferred from the insoluble PET particles to the aqueous bulk, regardless of whether it is pure aqueous T or part of a soluble fragment. We will call this the degree of solubilization, D^sol^, and it may be written:

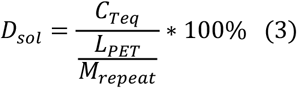

We note that a PET sample can be fully solubilized (D_sol_ ∼ 100%) even at moderate degrees of hydrolysis, D_hydrol_, because soluble fragments can retain a significant fraction of the ester bonds. Results in Fig. 2B (right ordinate) showed that towards the end of the 3 h experiments, all four enzymes had solubilized practically all PET (D_sol_ > 90%). For the two most efficient enzymes, IsP and LCC, these final hydrolysates contained fewer preserved ester bonds (D_hydrol_ ∼70%, Fig 2A) compared to HiC and TfC (D_hydrol_ ∼50-60%), but these differences were not large.

As mentioned above, direct comparisons of proton release and C_teq_ may provide further insights including information on endo-*vs*. exo-lytic modes of action. However, as D_hydrol_ changed rapidly in the initial stage (particularly for LCC) this approach requires a higher sampling rate than the one applied in Figs. 1-2. Hence, we conducted another series of measurements for LCC and HiC, which highlighted the initial stage of the reaction (5 s sampling intervals). These progress curves may be found in Fig. S1. The results are presented in a time-independent fashion in Fig. 3. Specifically, this figure shows the soluble product concentration, C_teq_ plotted against proton release. It appeared that HiC initially follows the diagonal (slope = 1) and hence releases equivalent amounts of protons and soluble T-equivalents. Later in the process, when [H^+^] and C_teq_ had reached around 70 µM, this changed to a mode with a higher production of soluble products (slope >1 in Fig. 3). By contrast, results for LCC fell well below the diagonal in the early part of the process, and we conclude that at least up to [H^+^]∼250 µM (corresponding to D_hydrol_=15%), LCC frequently cleaved bonds without a concomitant increment of T-equivalents in the bulk.

**Figure 3.**
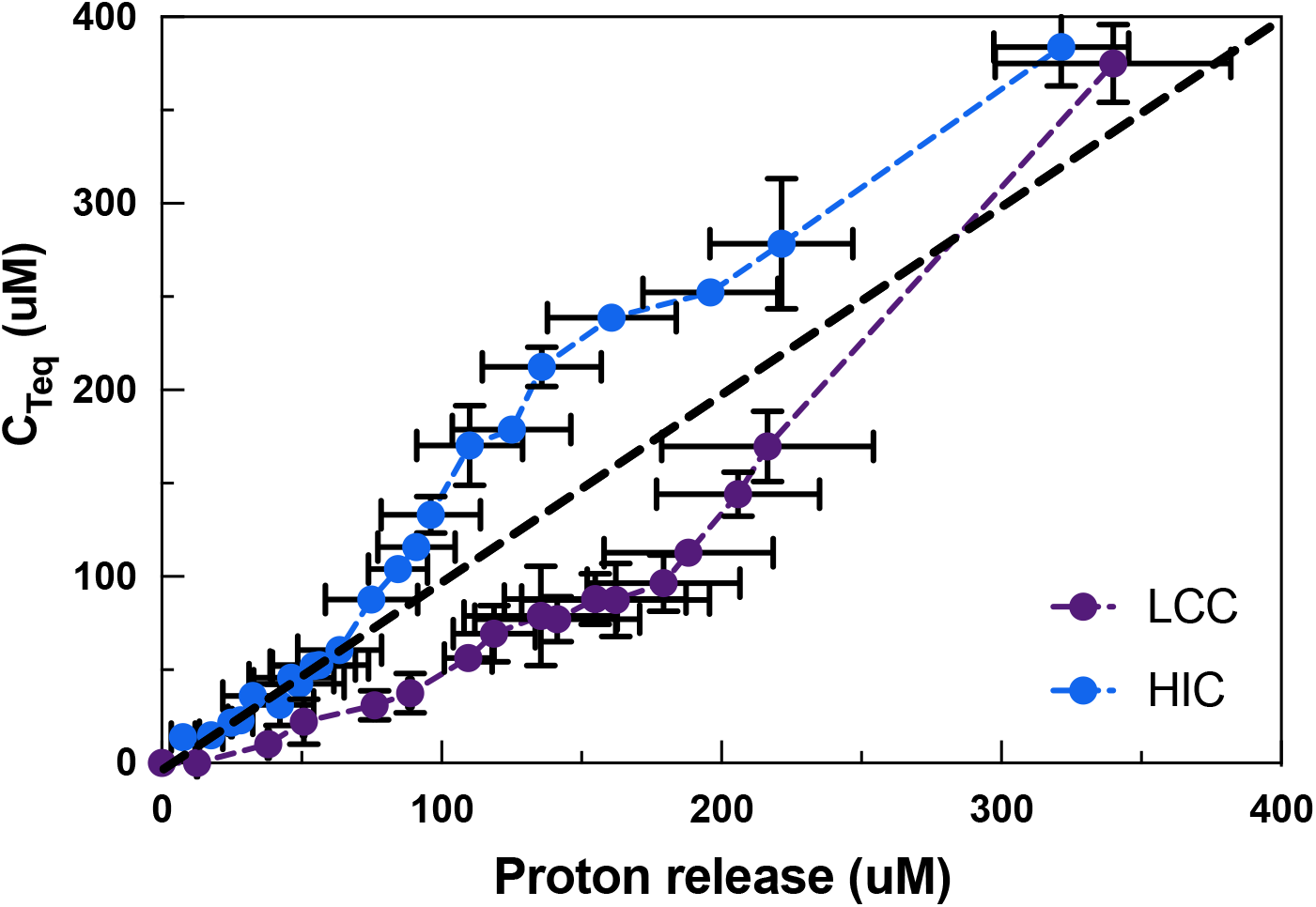
Total concentration of soluble fragments in terephthalic acid equivalents, C_teq_ (see Eq. 2) plotted as a function of proton release. The results are derived from progress curves measured by respectively pH-stat and RP-HPLC. Error bars represent the standard deviation of duplicate measurements.

### Steady-state kinetics for PET fragment hydrolysis

To assess the rate of bulk reactions in PET hydrolysis, we studied hydrolysis of the soluble fragments ET, ETE, TETE, and TETET by a standard Michaelis-Menten (MM) approach. Thus, we measured the initial steady-state rate, v^0^, and plotted it against the substrate concentration, [S]. We used non-linear regression of the MM equation:

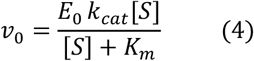

to derive values of the usual kinetic parameters k^cat^ and K^M^ (see Tab. S1 in the SI). Specificity constants, η=k^cat^/K^M^, calculated from the parameters in Tab. S1 are listed in Tab. 1. These results showed that ET was a very poor substrate for all investigated enzymes. The compound with one additional ethylene glycol moiety (ETE) was a much better substrate with specificity constants in the order of 10^3^ M^-1^s^-1^, and fragments with two or three aromatic rings showed markedly higher specificity constants in the range of 10^5^ M^-1^s^-1^. Interestingly, all specificity constants for the longest investigated fragment, TETET, fell within a factor of 2, and we conclude that the investigated enzymes showed a comparable and very high specificity for this substrate. Finally, it is of interest to note that the specificity constants consistently increased considerably when going from ETE to TETE and moderately between TETE and TETET. This would be in line with a mode of interaction where the substrate binds to subsites of at least two repeating units in PET.

### Kinetic modeling

We initially attempted deterministic modeling of PET hydrolysis, based on mass-action rate laws for potential reaction steps, but we found that this approach became overly complex, and as an alternative, we implemented a stochastic model. We considered random, iterative degradation of a polymer ([TE]_n_T) where n=200. This dimension of the polymer was chosen to reflect reported approximations of the molecular weight of PET^22–24^. We assigned a probability of bond breakage in iterative steps and conducted consecutive rounds until the polymer was mostly degraded. Guided by the experimental observations, we defined TETETE and smaller fragments as soluble, while ETETETE and longer fragments were considered insoluble. The probabilities of bond breakage in soluble fragments were scaled with the experimental specificity constants in Tab. 1. For insoluble substrates, the model distinguished between two types of enzyme reactions, endo-lytic and exo-lytic, with separate probabilities of bond breakage. We operationally defined exo-lytic reactions as those releasing soluble products (i.e. TETETE or shorter), while endo-lytic reactions did not. Finally, we noted in experiments that the smallest PET fragments, particularly ET, showed a tendency of non-enzymatic hydrolysis (autohydrolysis) at a rate that was comparable to or faster than the (very slow) enzymatic turnover of this molecule (c.f. Tab. 1). To account for this, we introduced a probability of ET autohydrolysis in the model. This value was fixed and based on experimental measurements of ET autohydrolysis (see Fig. S1 for experimental autohydrolysis results).

These criteria, which are summarized in Fig. 4, led to a stochastic model with 3 free parameters, namely the probability of respectively exo-lytic-(p_exo_), endo-lytic-(p_endo_) cuts in insoluble PET and an overall probability of bulk activity (p_solu_). A fourth, fixed parameter was used for the probability of an autohydrolysis event occurring (p_auto_). We used this model to interpret the experimental data in Figs. 1 and 2. Specifically, we performed regression to find the parameters, p_exo_, p_endo_, p_solu_, that best reproduced the progress curves for the five types of products listed in Fig. 1. Some examples of modeling results for HiC and LCC are shown in Fig. 5, and modeling results for IsP and TfC may be found in Fig. S3 in the supplementary section. Overall, it appeared that the three-parameter stochastic approach was able to account well for the experimental observations (Fig. 5, panels A and B). We used this to trace the formation and decay of different intermediates through the iterative steps, and hence elucidate the reaction pathway from polymeric PET to soluble monomers and fragments. Some results of this analysis for LCC and HiC are also presented in Fig. 5, panels C-F. It appeared from panels E and F that over the entire process, the average occurrence of the different modes of action defined in the model (see Fig. 4) was similar for LCC and HiC. Hence, hydrolysis of fragments in the bulk phase was the most common reaction (∼35% of all cleaved bonds), while the endo-lytic mode was the most common reaction on the insoluble substrate (∼30% of all cleaved bonds). Interestingly, autohydrolysis of ET was a significant decay route and accounted for over 15% of all esterolytic events in both systems (Fig. 5 E-F). While these values, accounting for the whole process, were similar, we saw distinct differences between the progress for LCC and HiC. For example, LCC showed a high initial level (panel C) of both endo-lytic cleavages in insoluble PET and bulk hydrolysis of soluble fragments. By contrast, HiC displayed a low initial endo-activity (Panel D), as well as a later maximum (compared to LCC) in the turnover rate of soluble fragments. Hence, the highest bulk activity of HiC occurred at around 35 min). This difference is also illustrated directly by the model parameters as p_endo_ was almost ten-fold higher for LCC compared to HiC. LCC also exhibited the highest p_exo_, but the difference in this parameter was moderate (a factor of 1.6, Tab. S2).

**Figure 4.**
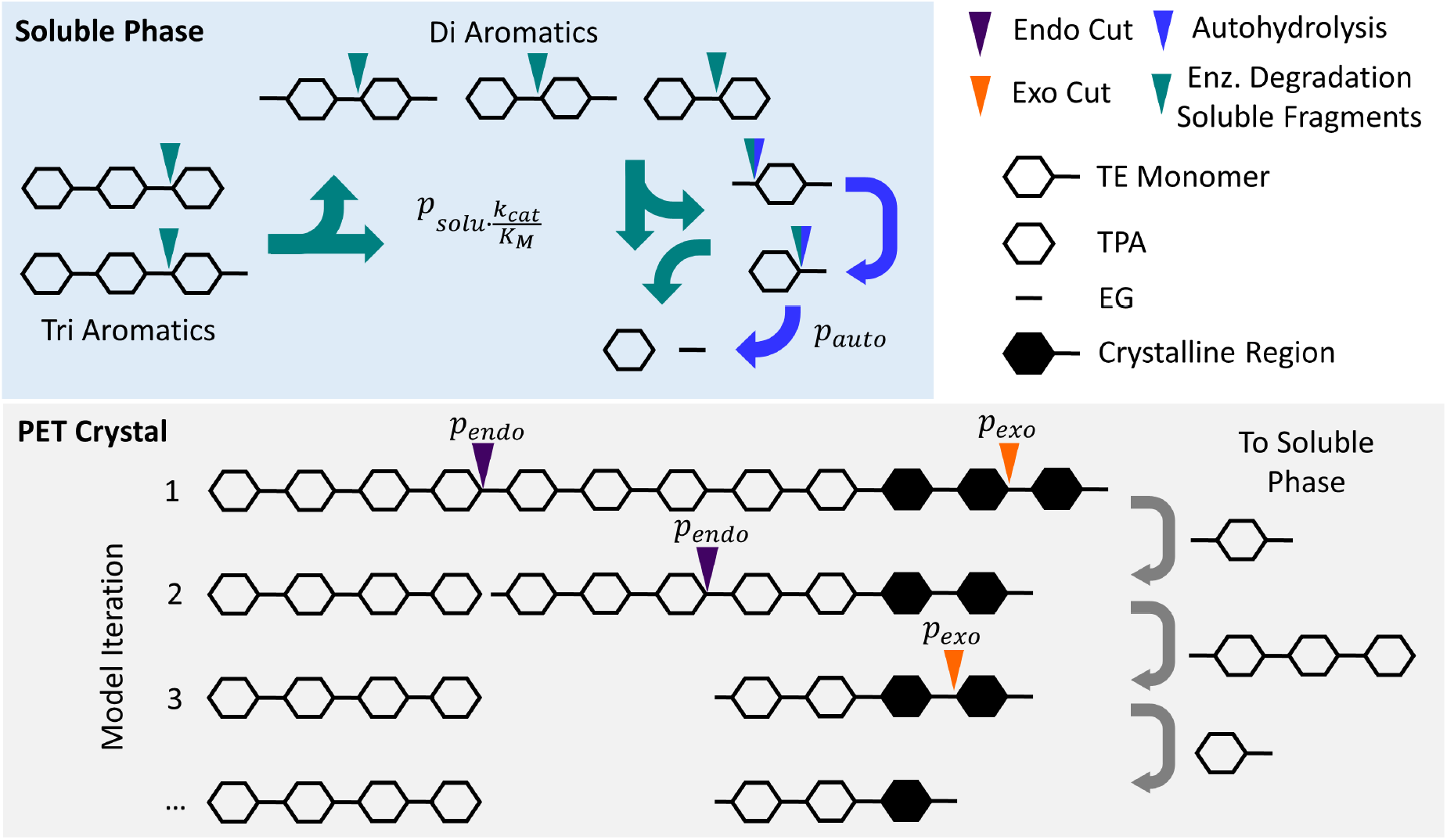
Overview of the stochastic models. Four different types of enzyme reactions were dictated by probabilities: For the insoluble substrate, the model differentiated between endo-(p_endo_) and exo-lytic activity (p_exo_). A small fraction of the insoluble substrate was defined as crystalline and was not susceptible to endo-lytic activity. In the bulk phase, we defined an overall probability of activity on soluble fragments (p_solu_) and scaled the event respectively to experimentally determined specificities of the enzymes for different PET fragments (see Tab. 1). A fourth parameter was defined for the probability of an autohydrolysis event occurring (p_auto_). The latter event was treated as a global fixed parameter and was not fitted to each enzyme individually as was the case for the remaining three types of enzymatic events. The model executed the described bond breakage over iterative consecutive rounds until the insoluble polymer was mostly converted to soluble monomers.

**Figure 5.**
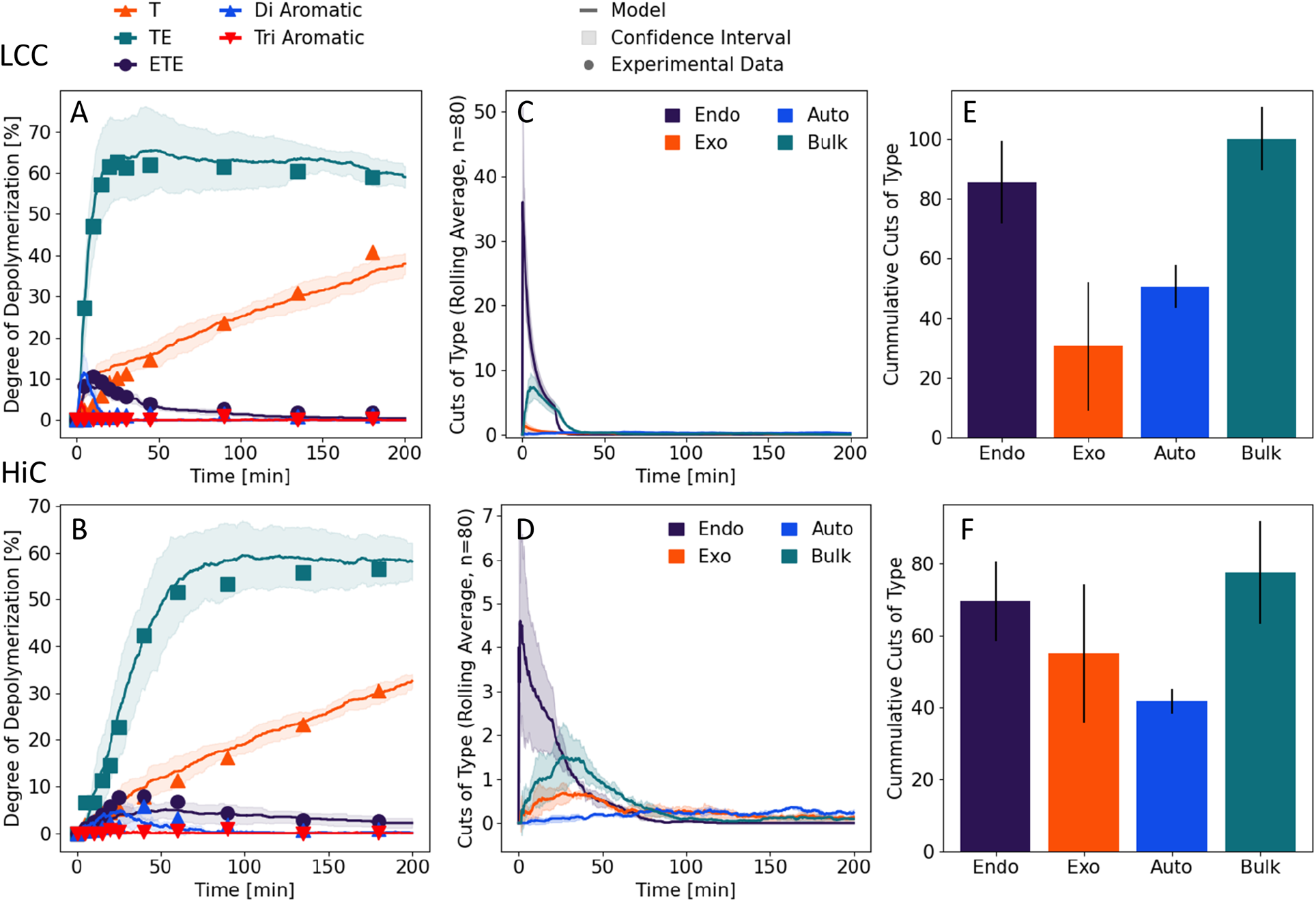
Results for the stochastic model fitted to experimental results for LCC (top panels) and HiC (lower panels). Average values and standard deviation of 5 repetitions of the stochastic models are shown. A/B) Values corresponding to terephthalic acid equivalents C_teq_ (see Eq. 2) occurring in each species during the reaction. C/D) Mode of hydrolytic event occurring over the time course of the stochastic model. E/F) Cumulative number of cuts per type.

The reaction pathways for LCC derived from the stochastic models are illustrated in a so-called Sankey plot in Fig. 6 (and Fig. S4 for HiC). This illustrates that the most common degradation pathway involves solubilization of larger fragments. Over half of all TPA that eventually accumulates has passed through a di-aromatic intermediate and around one third has passed through a soluble tri-aromatic intermediate. We conclude that these intermediates are released frequently, but as they are very rapidly hydrolyzed in the bulk (Tab. 1), their concentrations remain low. Nevertheless, the dynamics of fragment turnover appears to be central for the overall conversion rate.

**Figure 6.**
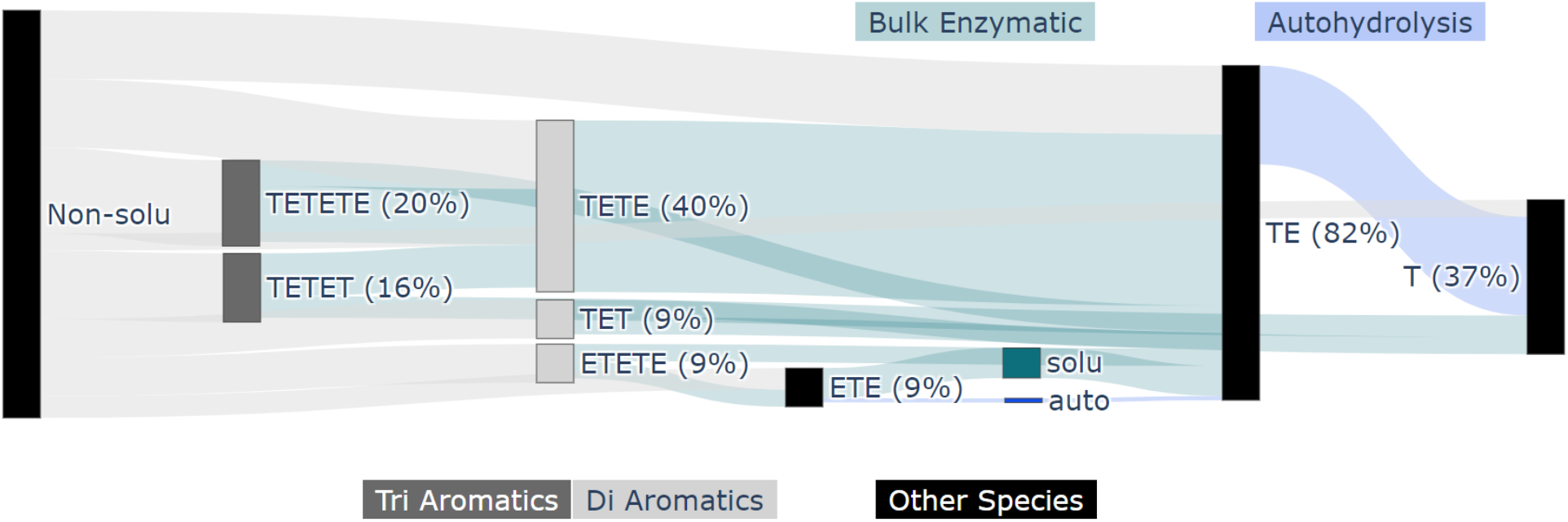
Sankey plot for LCC, illustrating the degradation pathway from insoluble PET towards monomers over the reaction course elucidated by the stochastic model. Different modes of action and soluble PET fragments are highlighted. The flux of terephthalic acid equivalents is scaled to the relative strength of the depicted linkage, and the percentages indicate the fraction of all terephthalic acid equivalents flowing through a given node. For the current discussion, it is important to note that soluble tri-aromatic (TETETE and TETET) and di-aromatic (TET, TETE and ETETE) compounds are very common intermediates. About 70% of all mono-aromatic products that eventually formed had gone through one of these intermediates.

## Discussion

Enzymatic degradation of insoluble polymers inherently involves a complex process, even in simple cases like the current, where an unbranched copolymer is broken down by a mono-component enzyme preparation. The complexity arises at least in part from the physical and chemical diversity of substrates, intermediates, and products that accumulate and decay during degradation. We propose that better insight into the pathways of interconversion between these compounds could help identify factors that promote or suppress the overall efficiency of the reaction. In the current work, we elucidated enzymatic degradation pathways for so-called nano-PET, which are amorphous particles with a size of about 140 nm^25 26^. This substrate makes stable aqueous suspensions and can be degraded within hours even at moderate enzyme dosages^17,25,27^, which provides a practicable platform for comparative analyses. Moreover, the amorphous nature provides an advantage by minimizing effects of crystallinity. This is important for comparative studies since several works have shown that even slight variations in substrate crystallinity plays a decisive role for the activity of PET degradability^11,23,28,29^.

In addition to measurements on nanoPET, we characterized the kinetics of PET-hydrolases on in-house synthesized, soluble PET fragments. These measurements led to two main conclusions, namely that PET fragments with two or three aromatic moieties were hydrolyzed very effectively and that the four investigated enzymes were quite similar with respect to their specificity for these soluble PET fragments (Tab. 1). These observations have some interesting corollaries. First, we note that specificity constants on TETET were in the range 3-4 × 10^5^ M^-1^s^-1^. This is notable inasmuch as it is comparable to (or even higher than) median values of specificity constants reported in broad meta-analyses of enzymes in general^30^ and hydrolases specifically acting on their native substrate^31^. It may be surprising that a non-native substrate is converted so efficiently, and the observation suggests that a PET structure can be readily positioned in a catalytically competent orientation in the active site of these enzymes. Possible misfits in the Michaelis complex have previously been discussed quite broadly for PET-hydrolases^32–35^. However, the high specificity constants demonstrated here suggest that this may not be a major barrier for PET hydrolysis, and points toward other aspects including the conformation and accessibility of the insoluble polymer as bottlenecks. As a second main aspect of the results in Table 1, we note that although the investigated enzymes are quite different with respect to their efficacy on polymeric PET^14,36^ (Fig. 2), their specificity constants on the tri-aromatic substrate TETET were quite similar. This observation further supports the view that the bottleneck for the hydrolysis of amorphous PET is not in the inner catalytic cycle, but associated to enzyme interactions with the solid substrate. In search of this bottleneck, we turn to the results for the insoluble PET.

**Table 1.** Specificity constants determined for HiC, TfC, IsP and LCC at an experimental temperature of 50 °C on soluble PET fragments. Standard deviations of duplicate measurements are shown in brackets.

| Substrate | HiC | Tfc | IsP | LCC |
| --- | --- | --- | --- | --- |
| | $\eta$ (M<sup>-1</sup>s<sup>-1</sup>) | $\eta$ (M<sup>-1</sup>s<sup>-1</sup>) | $\eta$ (M<sup>-1</sup>s<sup>-1</sup>) | $\eta$ (M<sup>-1</sup>s<sup>-1</sup>) |
| ET | ND<sup>a</sup> | 1.1<br>($\pm 3.5 \cdot 10^{-3}$) | ND | 40<br>($\pm 5.1$) |
| ETE | 5.5*10<sup>2</sup><br>($\pm 4.4 \cdot 10^1$) | 5.0*10<sup>3</sup><br>($\pm 1.6 \cdot 10^3$) | 3.3*10<sup>3</sup><br>($\pm 2.2 \cdot 10^2$) | 6.5*10<sup>3</sup><br>($\pm 1.3 \cdot 10^3$) |
| TETE | 3.7*10<sup>4</sup><br>($\pm 4.6 \cdot 10^3$) | 6.5*10<sup>4</sup><br>($\pm 6.8 \cdot 10^3$) | 9.9*10<sup>4</sup><br>($\pm 2.7 \cdot 10^4$) | 1.7*10<sup>5</sup><br>($\pm 5.1 \cdot 10^4$) |
| TETET | 2.7*10<sup>5</sup><br>($\pm 7.1 \cdot 10^4$) | 3.8*10<sup>5</sup><br>($\pm 9.8 \cdot 10^4$) | 4.0*10<sup>5</sup><br>($\pm 2.1 \cdot 10^5$) | 3.8*10<sup>5</sup><br>($\pm 2.5 \cdot 10^5$) |

a) ND = Not detected

Unlike LCC and IsP, the less proficient PET-hydrolases HiC and TfC showed a lag-phase both in HPLC (Fig. 2B) and pH-stat (Fig. 2A) measurements. One likely origin of this behavior appeared from Fig. 3, which showed the total concentration of dissolved terephthalic acid moieties (C_teq_, see Eq. 2) as a function of proton release. We note that this presentation disguises catalytic efficiency (there is no time dependence) and this provides some advantages for analyses of the reaction pathway. We found an initial slope for HiC which was essentially 1, and this implied that at low conversion, this enzyme released corresponding amounts of protons and soluble T-equivalents. This behavior is consistent with a reaction dominated by exo-lytic activity that primarily generates mono-aromatic products. For LCC, on the other hand, the initial slope was only about 0.5, and this suggested pronounced endo-lytic activity and conversion in the bulk (both giving proton release without a commensurate growth in dissolved T-equivalents). This conclusion was echoed in the model analysis, which demonstrated an initial endo-type activity for LCC that was almost an order of magnitude higher than for HiC (Fig. 5 C-D and Tab. S2). As a result, LCC promoted a substantial, early release of soluble PET fragments, which were converted quickly in the bulk (Fig. 5C). Model results for the other two enzymes, IsP and TfC, (Fig. S3 and Tab. S2) suggested that the reaction pathway of IsP resembled LCC while TfC was similar to HiC. Overall, this led to the conclusion that while all enzymes were comparably effective in converting fragments in the bulk phase, the more proficient PET-hydrolases excelled in rapid release of such fragments through endo-lytic activity. Moreover, slow release of fragments by HiC and TfC gave rise to a distinctive lag-phase (see Figs. 1-2).

Endo-lytic activity of PET hydrolases has been demonstrated by Eberl et al.^37^, who analyzed reaction products of TfC and a cutinase from *Fusarium solani* by mass spectrometry. Another study of this topic^38^ reported a positive correlation between the release of soluble products and increments in surface hydrophilicity of the PET-film. As endo-type activity generates carboxylic acid groups in the surface and hence increases hydrophilicity, this latter result is in line with our current observations. Several later works have confirmed the ability of different PET-hydrolases to conduct endo-type cuts (see Kawai et al. ^39^ for a review), and one particularly lucid example is the study by Zimmermann and co-workers^18^. This work used NMR to quantify endo-activity and concluded that the enzyme reaction started with endo-type chain scissions. This initial mode of action facilitated subsequent hydrolysis of neighboring ester bonds in an exo-type process. This overall interpretation again matches the current conclusions, although we propose that the most direct advantage of endo-lytic reactions was production of soluble fragments rather than chain-ends on the substrate surface.

## Conclusions

In conclusion, we have elucidated the reaction pathways for the degradation of amorphous nanoPET by four PET-hydrolases. In accordance with several other studies, our results showed that hydrolyzates were dominated by the small products T and ET, but we also observed larger di- and tri-aromatic, soluble PET fragments in low to moderate concentrations. Despite the low steady-state occurrence of these larger fragments, we propose that they are crucial for the overall degradation process. Thus, model analyses for LCC suggested that the majority (> 70%) of small products (T and ET) had been made via larger soluble intermediate (Fig. 6). These soluble fragments with two or three aromatic rings are hydrolyzed very efficiently in the aqueous bulk (Tab.1), and it follows that their steady state concentrations will remain low even if they are amply produced. In accordance with this interpretation, we found that endo-active enzymes quickly produced fragments and hence showed high overall activity against nanoPET. By contrast, enzymes with low endo-activity had a distinctive lag-phase and moderate overall efficacy. Ultimately, our work helps to pinpoint bottlenecks in the enzymatic degradation pathway of PET and serves to aid future works towards more efficient implementation of PET-hydrolases.

## Experimental procedures

### Enzymes

In this work, three cutinases were used, *Thermobifida fusca* (TfC) [GENBANK: AAZ54921.1]^40^, *Humicola insolens* (HiC) [GENBANK: AAE13316.1]^41^ and the leaf-branch compost cutinase (LCC) [GENBANK: AAE13316.1]^9^. In addition, we included a thermostable variant of the PETase (EC 3.1.1.101) from *Ideonella sakaiensis* (IsP), originally described by Austin et al^42^. All enzymes were expressed heterologous and purified as previously described^14,36^. Enzyme concentrations were determined by UV absorption at 280 nm using theoretical extinction coefficients.

### Substrates and chemicals

NanoPET was synthesized from PET-film with a crystallinity of approximately 5% (Goodfellow GmbH, Bad Nauheim, Germany, product number 991-473-70) using a precipitation and solvent evaporation procedure as described elsewhere^26^. The stock concentration of the suspended nanoPET was estimated to 1.66 g/L by dry-weight measurements in triplicates and the size of the final particles was determined by dynamic light scattering (137 ± 4.7 nm) using a Zetasizer Nano ZS (Malvern Panalytical, United Kingdom). Several compounds were used as substrates and/or standard samples for RP-HPLC analysis. T and ETE were purchased from Sigma-Aldrich, while ET, and the di- and tri-aromatic fragments (TET, TETE and TETET) were synthesized in-house starting from the mono-*t*Bu ester of terephthalic acid (*t*Bu-T-OH, CAS 20576-82-3). *t*Bu-T-OH was purchased from Enamine Ltd (NJ, USA). Alternatively, it can be synthesized according to the procedure by Mineno et al.^43^. In contrast to previous Bn protecting group strategies^44^, the *t*Bu-group improves solubility of the intermediates in organic solvents like CHCl^3^ and is easily removed under acidic conditions. Synthesis of ETETETE was inspired by Boutevin et al.^45^. The products were characterized by NMR and liquid chromatography mass spectrometry (LC-MS) analysis, showing >95% purity.

### Progress curves

Samples of 4 mL, 166 mg/L nanoPET dispensed in water and with pH adjusted to 8.0 by 50 mM NaOH were loaded into a temperature-controlled reactor and covered by an air-tight lid. The sample was allowed to thermally equilibrate during magnetic stirring at 200 rpm and this stirring was continued throughout the experiment. The reaction was started by the injection of enzyme to a final concentration of 0.30 µM (LCC, HiC and TfC) or 0.20 µM (IsP), and progress curves were monitored over 3 h. The experimental temperature was 50 °C for LCC, HiC, and TfC. IsP is less thermostable, and this dictated a lower experimental temperature of 40°C. All progress curves were measured in duplicates.

Progress of the enzymatic reaction in these samples was monitored by two independent analytic methods, pH-stat and RP-HPLC. The pH-stat was built upon an open-source software template and was controlled by Arduino microcontrollers with a USB connection, as described previously^46^. Here, the pump module consisted of a syringe pump module responsible for delivering 50 mM NaOH to adjust the pH to 8.00. As described by Miksch et al.^47^ a typical course of NaOH consumption over time was divided into different stages. Initially, the pH of the suspension was adjusted to pH 8.00. Subsequently, enzyme was added and dosing of small quantities of NaOH was monitored that were prompted by unspecific reactions in the suspension (enzyme blank). Finally, nanoPET was added and NaOH was continuously delivered to adjust for the release of protons from esterolytic activity and unspecific reactions.

For RP-HPLC analysis we regularly withdrew 10 small subsets of 60 µL of the reacting solution. In the initial phase of the reaction, where activity was most extensive, samples were withdrawn every 5 min. All samples withdrawn were quenched by the addition of 3.5 µL of 5 M HCl. Subsequently, the content of soluble PET fragments in the samples was analyzed by RP-HPLC (Hewlett Packard, ChemStation 1100 series: Capital HPLC (250 mm x 4.6 mm) C18 column). Samples were injected in volumes of 20 µL using 7.5 mM Formic acid and 5% v/v acetonitrile at a ratio of 1:4 for 25 min and eluted by acetonitrile. The flow rate was kept at 0.5 ml/min at 40 °C. We used UV detection at 240 nm and peak analysis was performed in the Agilent ChemStation LC 3D software. The results were quantified against standards of T, ET, ETE, TET, TETE, and TETET. Duplicates and substrate blanks (for quantification of autohydrolysis) were included for all reactions.

For HiC and LCC, an additional experiment was conducted focusing on the initial stage of the reaction. For this purpose, the enzyme concentration was reduced to 0.18 µM and subsets were retrieved more frequently (every 5 s and 20 s over the first two minutes for LCC and HiC, respectively). To allow for more subsets, the reaction volume was increased to 10 mL and 20 subsets of 60 µL were withdrawn from the reaction solution of both enzymes.

### Kinetics on soluble PET fragments

Activity on the PET fragments ET, ETE, TETE, and TETET was assayed in low binding microplates (Greiner Bio-One™ 655900). Reactions were performed in duplicates in 250 µL volumes with substrate concentrations varying from 0 and 2 mM. The plates were incubated in an Eppendorf thermomixer at 50°C, 1100 rpm with contact times of 10 min (ETE, TETE, TETET, and ETETETE) or 2 hours (ET). Due to the short extend of the reaction time, the less thermostable IsP was also assayed at 50°C. Enzyme concentrations were kept low to avoid secondary reaction products and ranged from 0.01-0.5 µM, depending on substrate and enzyme. Duplicates and substrate blanks (for quantification of autohydrolysis) were included, and all reactions were quenched by addition of HCl and analyzed by RP-HPLC as described above.

### Stochastic Model

An iterative stochastic model with four parameters (three free, one fixed) was designed to elucidate the degradation pathways of PET for the four PET-hydrolases (see Fig. 4). The four parameters included the probabilities per minute to perform: i + ii) endo- and exo-cuts on the insoluble part of the substrate iii) enzymatic hydrolysis of already solubilized fragments in the bulk phase iv) autohydrolysis of ET and ETE. The probability for degrading solubilized fragments was weighted correspondingly with the specificity of the enzyme for the respective fragment (see Tab. 1).

The system was initialized with a single chain of PET consisting of 200 monomers with terephthalic acids on both ends (e.g., ([TE]_n_T where n=200). Crystallinity was assumed for 1% of the monomers and disallowed any endo-cuts in a proximity of ±5 monomers. Additionally, endo-cuts were prohibited within two monomers of existing ends. Contrarily, exo-cuts were permitted to cleave off any fragment consisting of an aromatic moiety (i.e., T, ET or ETE). PET fragments of the size of TETETE and smaller were assumed soluble and thus accessible for further degradation in the bulk phase (see Fig 4).

The model executed the defined events of bond breakage over consecutive iterative rounds until the insoluble polymer was predominantly converted to soluble monomers. Meanwhile, progress curves for the accumulation and decay of substrates liberated from the insoluble PET chain were monitored. It was arbitrarily chosen that four iterations of the stochastic model corresponded to one minute of our experimental data shown in Figs. 1 and 2. In total, the models were run for 800 iterations (reflecting 200 minutes of the experimental data). Models were run in repetitions of 5. Differential evolution was used to fit the models (average T-equivalents (C_teq_, see Eq. 2) in each species at the corresponding iteration) to the experimental product profiles (T-equivalents in each species) in a least-squares fashion.

In an initial round, the probability of autohydrolysis was estimated through fitting and then fixed to the average value of all enzyme systems (p_auto_=0.19%). Subsequently, only the probability of exo-cuts (p_exo_), endo-cuts (p_endo_), and degradation of solubilized products (p_solu_) were fitted for each enzyme system. The model was implemented and visualized in python 3.8.3^46^ with scipy 1.5.0^48^ and plotly 5.5.0^49^.

## Supporting information

Full supporting information

## Data availability

All data that support the findings of this study are included in the published article and its Supporting information file.

## Acknowledgements

The authors acknowledge Josu Arano Noriega for technical assistance with laboratory experiments.

## Abbreviations

ETE: Bis(2-hydroxyethyl) terephthalate;
E: Ethylene glycol;
LC-MS: liquid chromatography mass spectrometry;
MM: Michaelis-Menten;
PET: Poly(ethylene terephthalate);
ET: Mono(2-hydroxyethyl) terephthalate;
RP-HPLC: Reversed-phase high-performance liquid chromatography;
MALDI-TOF: matrix-assisted laser desorption/ ionization time-of-flight;
NMR: Nuclear magnetic resonance;
T: Terephthalic acid:

## Table of contents

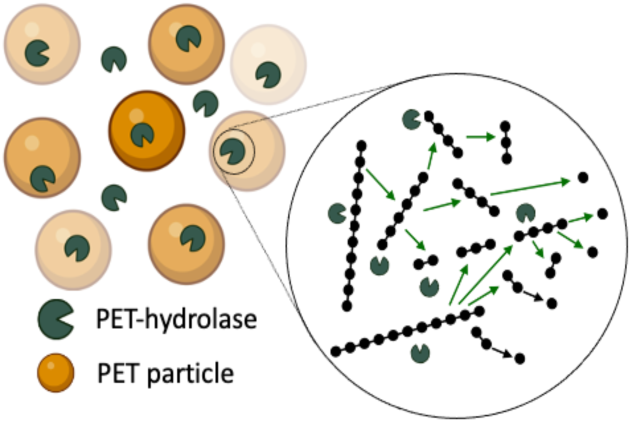
Enzymatic degradation of PET. Here, we present their reaction pathway of four PET-hydrolases. Moreover, we monitor the accumulation and decay a various released PET fragments and determine their specificity of the enzymes for these products. Specific modes of enzymatic reactions were elucidated using a stochastic model.

## Footnotes

1 mono-(2-hydroxyethyl) terephthalate and bis-(2-hydroxyethyl) terephthalate are often abbreviated MHET and BHET respectively. However, in the current work we use several still longer fragments and hence apply the simpler abbreviation of listing the ethylene glycol (E) and terephthalate (T) moieties of the molecules. Mono-(2-hydroxyethyl) terephthalate, for example becomes ET, bis-(2-hydroxyethyl) terephthalate, ETE and so forth.

