## Supplementary material for "Reaction pathways for the enzymatic degradation of poly(ethylene terephthalate): What characterizes an efficient PET-hydrolase?": Full supporting information

### SUPPLEMENTARY INFORMATION

1. **Table S1.** *Steady state kinetics on soluble reaction fragments*
2. ***Figure S1.*** *Progression curves for the initial stage of the reaction*
3. ***Table S2.*** *Parameters for the stochastic model derived by differential evolution.*
4. ***Figure S2.*** *Rate of autohydrolysis of ETE  and ET*
5. ***Figure S3.*** *Results for the stochastic models fitted to experimental results for LCC and HiC*
6. ***Figure S4***. *Exemplary Sankey plot for HiC*

**Table S1.** *Steady state kinetics on soluble reaction fragments. $k_{cat}$ and $K_M$ values were determined for HiC, TfC, IsP and LCC at an experimental temperature of 50 ˚C. Standard deviations of duplicate measurements are shown in brackets.*

| | HiC | | TfC | | IsP | | LCC | |
|---|---|---|---|---|---|---|---|---|
| **Substrate** | $K_M$ (mM) | $k_{cat}$ ($s^{-1}$) | $K_M$ (mM) | $k_{cat}$ ($s^{-1}$) | $K_M$ (mM) | $k_{cat}$ ($s^{-1}$) | $K_M$ (mM) | $k_{cat}$ ($s^{-1}$) |
| ET | **ND**[a] | **ND** | **3.8**[b] (± 0.091) | **0.043** (± 8.5E-4) | **ND** | **ND** | **1.2**[b] (± 0.13) | **0.048** (± 0.0033) |
| ETE | **3.7** [b*] (± 0.24) | **2.0** (± 0.092) | **0.23** (± 0.067) | **1.1** (± 0.12) | **1.4** [b*] (± 0.082) | **4.8** (± 0.15) | **0.96** [b] (± 0.16) | **6.2** (± 0.62) |
| TETE | **0.077** (± 0.0091) | **2.9** (± 0.098) | **0.17** (± 0.016) | **11** (± 0.35) | **0.14** (± 0.036) | **14** (± 1.3) | **0.12** (± 0.035) | **21** (± 1.7) |
| TETET | **0.089** (± 0.021) | **24** (± 2.9) | **0.17** (± 0.038) | **62** (± 7.3) | **0.058** (± 0.029) | **23** (± 4.2) | **0.79**[b] (± 0.41) | **300** (± 122) |

a) ND = Not detected
b) Not saturated

*Data from Arnling Bååth et al. 2020 (ChembioChem)

#### Initial stage of the reaction

Fig. 3 in the main section shows the total concentration of dissolved $C_{Teq}$ (see Eq. 2) as a function of proton release. This representation of the data disguises the enzyme efficiency by not reflecting any given time dependency. The figure below shows the progress curves for both $C_{Teq}$ and the release of protons as function of time.

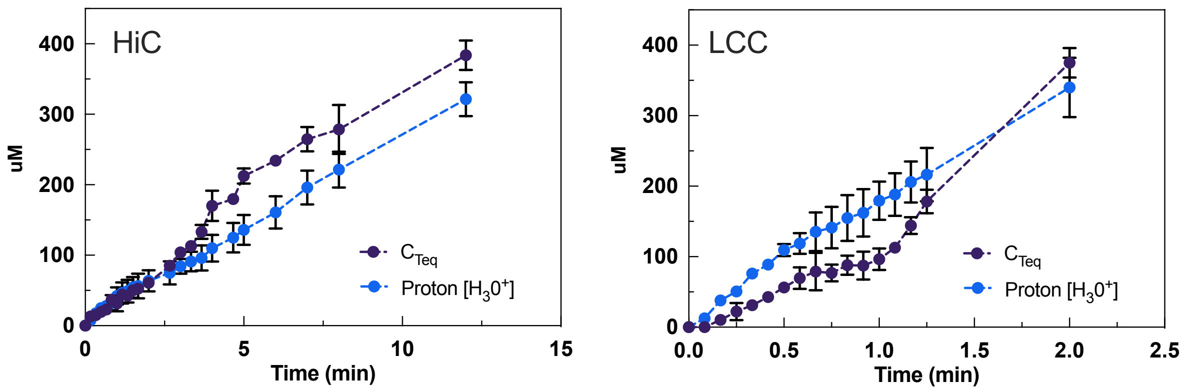


***Figure S1.*** *Progression curves for the initial stage of the reaction for both terephthalic acid equivalents, $C_{Teq}$ (see Eq. 2) and release of protons plotted as a function of time. The results are derived from measurements by respectively pH-stat and RP-HPLC. Error bars represent the standard deviation of duplicate measurements.*

***Table S2.*** *Parameters for the stochastic model derived by differential evolution. The parameters are probabilities per minute to perform a certain type of esterolytic activity. Endo-lytic ($p_{endo}$) and exo-lytic ($p_{exo}$) activities on the insoluble substrate as well as enzymatic hydrolysis of solubilized fragments and autohydrolysis of ETE and TE fragments were distinguished (see Fig. 4). The probability of the enzyme to act on soluble fragments ($p_{solu}$) is weighted with the specificity constant (see Tab. 1), which varies for each enzyme. The probability of autohydrolysis ($p_{auto}$) was treated as a global parameter and not fitted to each enzyme individually.*

| **Enzyme** | $p_{endo}$ [-] | $p_{exo}$ [-] | $p_{solu}$ [-] | $p_{auto}$ [-] |
|---|---|---|---|---|
| HiC | 1.2% | 0.7% | 0.31% | 0.19% |
| LCC | 12.5% | 1.1% | 0.12% | 0.19% |
| TfC | 0.6% | 0.9% | 0.12% | 0.19% |
| IsP | 6.6% | 0.7% | 0.11% | 0.19% |

**Autohydrolysis of MHET of BHET**

While developing the stochastic model intended to examine the enzymatic degradation pathway of PET towards soluble monomers, we obtained results suggesting that the contribution from enzymatic activity alone, could not account for the presence of TPA. Hence, an experiment was conducted to investigate the rate of autohydrolysis of ET and ETE under the conditions used to obtain experimental results for the enzymatic degradation course of PET (see Figs. 1-2). The autohydrolysis of ET and ETE was examined for two initial concentrations that were in proximity to the amount of the respective product detected during our experimental runs (see Fig. 1).

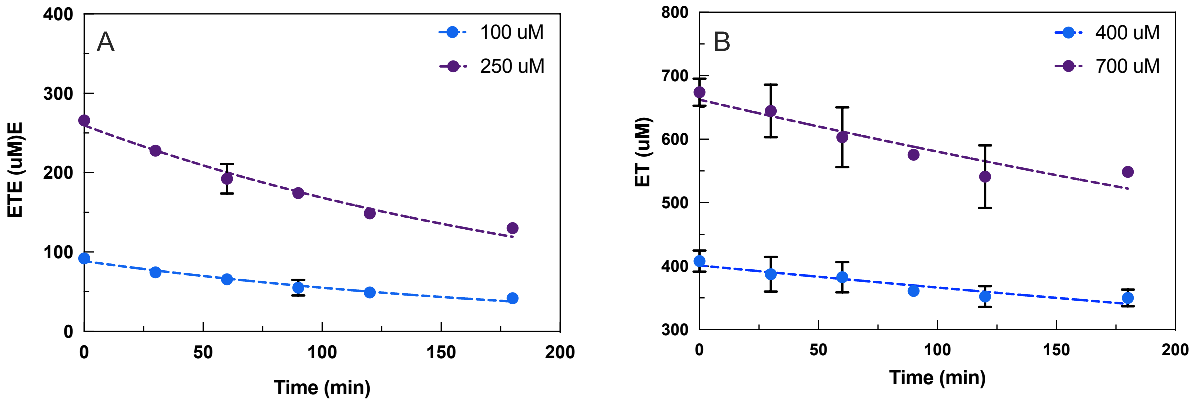


***Figure S2.*** *Rate of autohydrolysis of ETE (A) and ET (B) examined under the same experimental conditions used to obtain the results shown in Figs. 1 and 2. RP-HPLC analysis was used to detect the presence of either ET and ETE and the autohydrolysis of the respective product was tracked over time. Based on our results, the half-life of ETE and ET was estimated to 153.0 ± 10.3 min and 643 ± 168.3 min, respectively.*

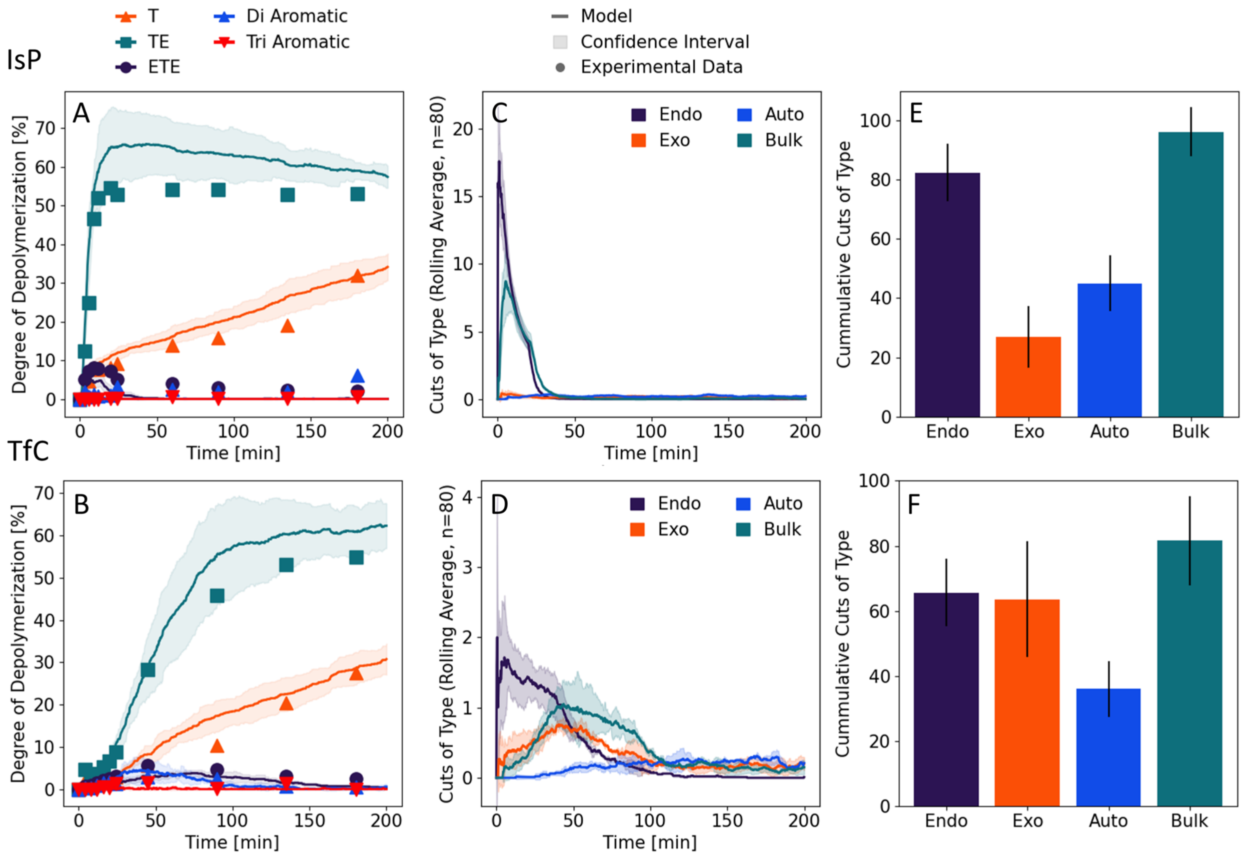


***Figure S3.*** *Results for the stochastic models fitted to experimental results for LCC (top panels) and HiC (lower panels). Average values and standard deviation of 5 repetitions of the stochastic models are shown. A/B) Value corresponding to terephthalic acid equivalents* $C_{Teq}$ *(see Eq. 2) occurring in each species during the reaction. C/D) Mode of hydrolytic event occurring over the time course of the stochastic model. E/F) Cumulative amount of cuts per type.*

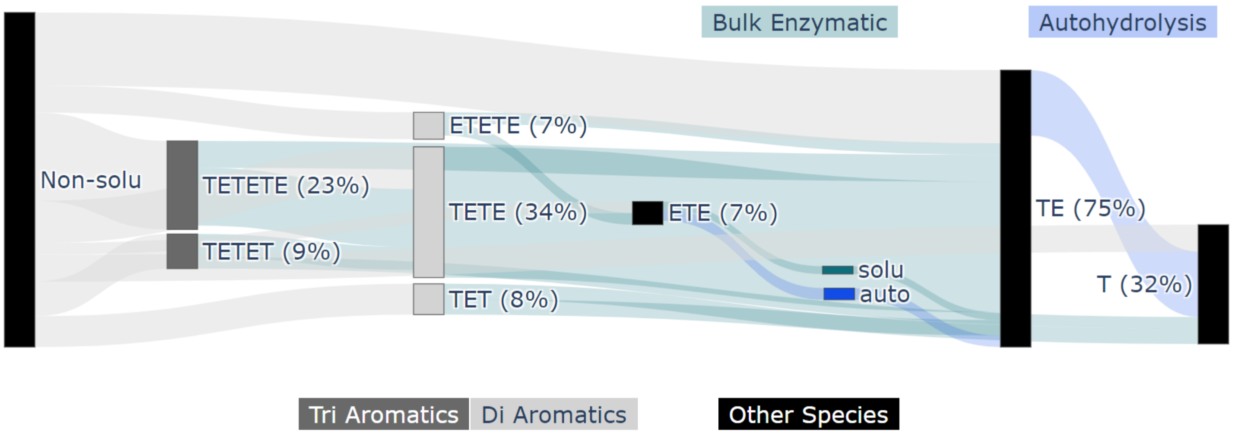


***Figure S4***. *Exemplary Sankey plot for HiC depicting the degradation pathway from insoluble PET towards monomers over the reaction course elucidated by the stochastic model. Different modes of action and soluble PET fragments are highlighted. The flux of terephthalic acid equivalents ($C_{Teq}$, see Eq. 2) is scaled to the relative strength of the depicted linkage, and the percentage of the total $C_{Teq}$ flowing through a node is highlighted.*
